# Global brain signal in awake rats

**DOI:** 10.1101/802884

**Authors:** Yuncong Ma, Zilu Ma, Zhifeng Liang, Thomas Neuberger, Nanyin Zhang

## Abstract

Although often used as a nuisance in resting-state functional magnetic resonance imaging (rsfMRI), the global brain signal in humans and anesthetized animals has important neural basis. However, our knowledge of the global signal in awake rodents is sparse. To bridge this gap, we systematically analyzed rsfMRI data acquired with a conventional single-echo (SE) echo planar imaging (EPI) sequence in awake rats. The spatial pattern of rsfMRI frames during peaks of the global signal exhibited prominent co-activations in the thalamo-cortical and hippocampo-cortical networks, as well as in the basal forebrain, hinting that these neural networks might contribute to the global brain signal in awake rodents. To validate this concept, we acquired rsfMRI data using a multi-echo (ME) EPI sequence and removed non-neural components in the rsfMRI signal. Consistent co-activation patterns were obtained in extensively de-noised ME-rsfMRI data, corroborating the finding from SE-rsfMRI data. Furthermore, during rsfMRI experiments we simultaneously recorded neural spiking activities in the hippocampus using GCaMP-based fiber photometry. The hippocampal calcium activity exhibited significant correspondence with the global rsfMRI signal. These data collectively suggest that the global rsfMRI signal contains significant neural components that involve coordinated activities in the thalamo-cortical and hippocampo-cortical networks. These results provide important insight into the neural substrate of the global brain signal in awake rodents.

## Introduction

Compelling evidence suggests that the intrinsic brain activity plays an essential role in brain function. For instance, this spontaneously fluctuating brain activity consumes a major portion of brain’s energy budget (termed as brain’s dark energy (Raichle 2010)), much higher than that used for external tasks (Raichle 2006), and anomalies in intrinsic brain activity are tightly linked to brain disease (Zhang and Raichle 2010). The prominent utility of intrinsic brain activity, typically measured by resting state functional magnetic resonance imaging (rsfMRI), is to assess inter-areal resting-state functional connectivity (RSFC) (Biswal et al. 1995). Given its simplicity, whole-brain coverage and sensitivity, this method has been widely applied in human and animal studies and has revolutionized our understanding of brain network organization (Biswal et al. 2010; Smith et al. 2009; Liang et al. 2011).

An interesting research topic in intrinsic brain activity is the global brain signal, which is defined as the averaged rsfMRI signal across all brain voxels. The global signal was initially introduced as a nuisance to regress out in rsfMRI data preprocessing (Aguirre et al. 1997). The rationale underlying global signal regression is that distributed spontaneous neural activities are semi-random and out of phase, which will be cancelled out when rsfMRI signal is averaged across the whole brain. Therefore, the global rsfMRI signal can be treated as a nuisance dominated by non-neural fluctuations. Indeed, vascular signals from large veins show high temporal correlation to the global signal (Colenbier et al., 2019). As RSFC is assessed by temporal correlations between distributed rsfMRI signals, it is particularly susceptible to non-neural confounds like head motion (Van Dijk et al. 2012; Power et al. 2012; Power et al. 2015; Satterthwaite et al. 2012), respiration (Birn et al. 2008) and cardiac pulsation (Chang et al. 2009), which all affect the rsfMRI signal globally and can cause systematic bias in RSFC quantification (Liu 2016). Therefore, regressing out the global rsfMRI signal represents an effective method for removing these non-neural artifacts (Ciric et al. 2018), and has been widely used (Fox et al. 2005).

Despite its effectiveness, doubt has been casted on the rationale of global signal regression (Murphy et al. 2009) and it has been suggested that the global signal might have important neural components (Liu et al. 2018; Turchi et al. 2018; Scholvinck et al. 2010; Yang et al. 2014). Simultaneous recordings of electrophysiology and rsfMRI data revealed synchronized neural activity across the brain, which was also strongly correlated to brain-wide rsfMRI signals (Wen and Liu 2016; Scholvinck et al. 2010), showing that the global signal has neural basis. In addition, a series of studies reported that the global brain signal was tightly linked to the vigilance level in human subjects (Rack-Gomer and Liu 2012; Wong et al. 2012, 2013; Chang et al. 2016a; Liu et al. 2018), which suggests that the global signal plays an important functional role. Furthermore, altered global brain signal was reported in patients with schizophrenia, indicating that the global signal could be an endophenotype of psychiatric disorders (Yang et al. 2017; Yang et al. 2014). Taken together, these studies have greatly advanced our understanding of the neural basis and physiologic function of the global signal in humans.

Recent studies in anesthetized animals have also shed light on potential neural contributions to the global signal. Wide field imaging of calcium signals showed synchronized neural activation across a large proportion of the cortex (Ma et al. 2016; Matsui et al. 2016). In addition, brain’s co-activation patterns in mice demonstrated specific phase relationship to the global signal fluctuation (Gutierrez-Barragan et al., 2018). However, our knowledge of the global signal in awake rodents is still lacking. Investigating this issue in awake animals is important as it avoids the potential confounding effects of anesthesia on animals’ physiologic states and the global signal (Gao et al. 2016; Liang et al. 2012b; Liang et al. 2015a; Ma et al. 2017; Smith et al. 2017; Hamilton et al. 2017), and permits linking brain activity measured to behavior (Liang et al. 2014; Dopfel et al. 2019). To bridge this gap, here we systematically investigate the global brain signal using the awake rat fMRI approach established in our lab (Liang et al. 2011; Zhang et al. 2010; Perez et al. 2018; Dopfel and Zhang 2018). We first examine the spatial patterns of rsfMRI volumes during peaks of the global signal. Our data demonstrate that the global signal in awake rats might be linked to coordinated neural activities in the thalamo-cortical and hippocampo-cortical networks. To confirm this concept, we acquire rsfMRI data using a multi-echo (ME) echo planar imaging (EPI) sequence and use its advantage to remove non-neural components in rsfMRI signal (Kundu et al. 2013; Kundu et al. 2012; Kundu et al. 2014). Consistent co-activation patterns are obtained in extensively de-noised ME-rsfMRI data. Furthermore, during rsfMRI experiments we simultaneously record spiking activities in the hippocampus using GCaMP-based fiber photometry and find significant correspondence between hippocampal calcium signal and the global rsfMRI signal. These data collectively suggest that the global signal in awake rodents contains important neural components involving activities in the thalamo-cortical and hippocampo-cortical networks.

## Methods and Materials

### Animal Preparation

All procedures were conducted in accordance to approved protocols from the Institutional Animal Care and Use Committee (IACUC) of the Pennsylvania State University. 92 adult male Long-Evans rats (250-500g) were used in this study. 71 rats (133 scans) were scanned using single-echo rsfMRI (SE-rsfMRI), and 21 rats (43 scans) were scanned using ME-rsfMRI. Part of the SE-rsfMRI data were used in previous publications (Ma et al. 2018; Ma and Zhang 2018) and was reprocessed for the purpose of the present study. All animals were housed and maintained on a 12hr light: 12hr dark schedule, and provided access to food and water *ad libitum*.

To minimize stress and motion during imaging at the awake state, animals were acclimated to the scanning environment for 7 days (see details described in (Dopfel and Zhang 2018; Gao et al. 2016)). Briefly, rats were briefly anesthetized (5 min) under isoflurane (2-4%) and placed in a body and head restrainer. After this setup, isoflurane was discontinued, and animals were allowed to regain full consciousness. The restrainer with the animal was then placed in a mock MRI scanner where prerecorded sounds of MRI pulse sequences were played. The exposure time started from 15 min on day 1, and was incrementally increased by 15 min each day (days 2 and 3) up to 60 minutes (days 4, 5, 6 and 7). This setup mimicked the scanning condition inside the magnet. A similar acclimation approach has also been used by other research groups (Bergmann et al. 2016; Yoshida et al. 2016; Chang et al. 2016b).

### MRI Experiments

Using the method described in the acclimation procedure, rats were placed in an identical head restrainer with a built-in birdcage radiofrequency coil. Isoflurane was discontinued after the animal was set up. Imaging began ~30min after rats were placed in the scanner while they were fully awake. Image acquisition was performed at the High Field MRI Facility at the Pennsylvania State University on a 7T Bruker 70/30 BioSpec running ParaVision 6.0.1 (Bruker, Billerica, MA). After a localizer scan, T1-weighted structural images were acquired using a rapid imaging with refocused echoes (RARE) sequence with the following parameters: repetition time (TR) = 1500ms, echo time (TE) = 8ms, matrix size = 256 × 256, field of view (FOV) = 3.2 × 3.2 cm^2^, slice number = 20, slice thickness = 1mm, RARE factor = 8, and repetition number = 8. For the SE-rsfMRI experiment, one to three SE-rsfMRI scans were acquired using a single-shot gradient-echo EPI pulse sequence with the following parameters: TR = 1000ms, TE = 15ms, flip angle = 60°, matrix size = 64 x 64, FOV = 3.2 × 3.2 cm^2^ (in-plane resolution = 0.5 × 0.5mm^2^), slice number = 20, slice thickness = 1 mm. 600 SE-rsfMRI volumes (10 min) were acquired for each SE-rsfMRI scan.

21 animals (body weight = 307 ± 19 g) were scanned in an ME-rsfMRI experiment. The setup was the same as the SE-rsfMRI experiment. As the scanner noise was louder for the ME-EPI sequence, earplugs were used in rats to reduce acoustic noise during ME-rsfMRI data acquisition. For each animal, one to three ME-rsfMRI scans were acquired using a single-shot ME-EPI sequence (Fig. 1A) with TR = 1500 ms, TEs = 7.6, 16.4, 25.2 and 34.0 ms, flip angle = 75°, matrix size = 48 × 32, FOV = 2.4 × 1.6cm^2^ (in-plane resolution = 0.5 × 0.5 mm^2^), slice number = 18, slice thickness = 1 mm. 400 ME-rsfMRI volumes (10 min) were acquired for each ME-EPI scan.

**Figure 1.**
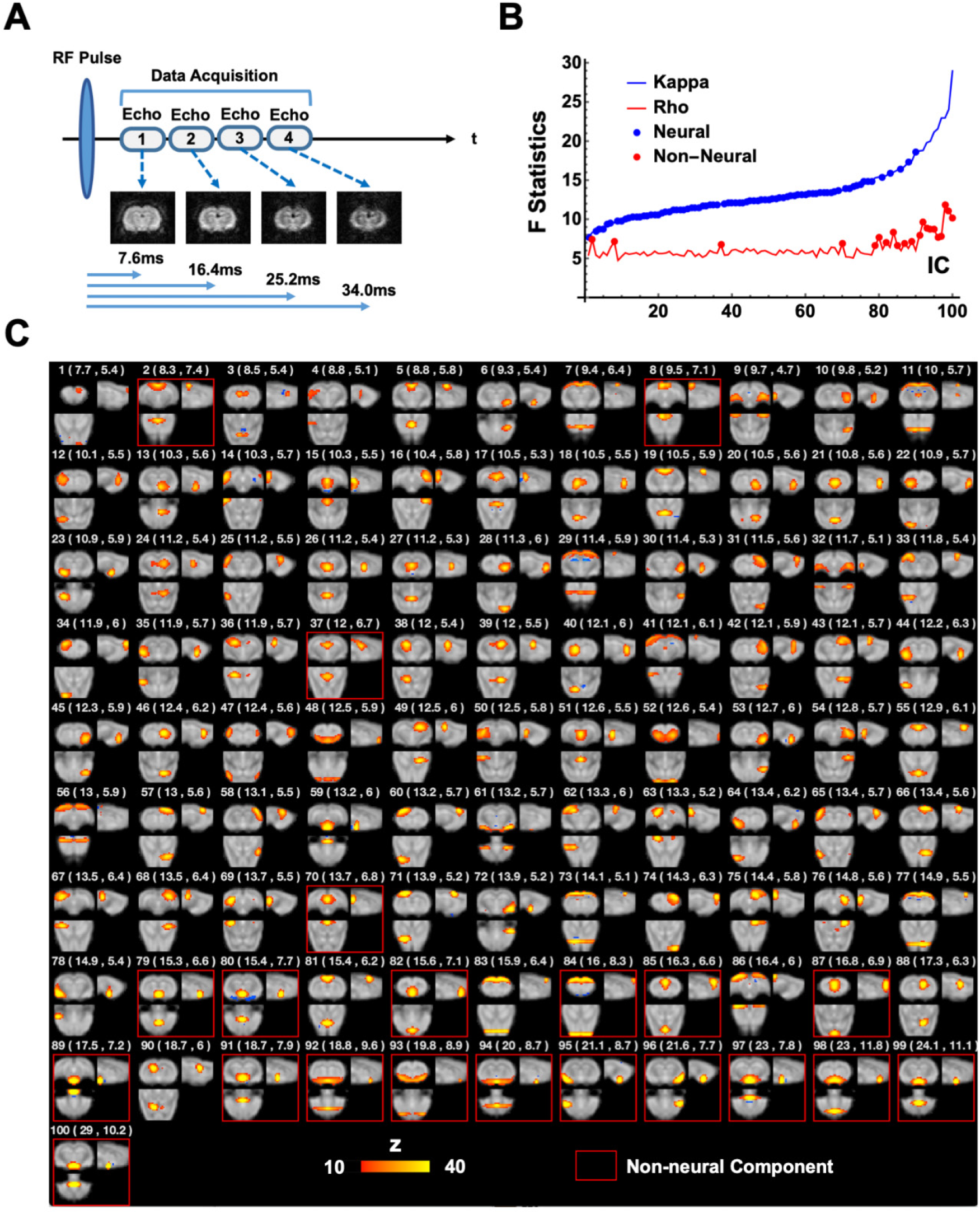
ME-EPI data acquisition and processing. (**A**) Diagram of the ME-EPI pulse sequence. Four images were acquired per rsfMRI volume, each at a different TE. (**B**) BOLD (κ) or non-BOLD (ρ) weights of individual ICA components (blue and red lines), sorted by their κ values. 79 neural components (blue dots) and 21 non-neural (red dots) were identified. (**C**) Spatial map, as well as *κ* and *ρ* values (listed in brackets) for each individual component. Non-neural components were highlighted by red boxes.

### Data Analysis

Analyses of SE-rsfMRI and ME-rsfMRI data were carried out using MATLAB 2017b (The MathWorks, Inc., Natick, MA). Mathematica 11.3 (Wolfram Research, Inc., Champaign, IL), MATLAB and ITK-SNAP (Yushkevich et al. 2006) were used for data visualization.

#### SE-rsfMRI data preprocessing

SE-rsfMRI data were preprocessed using the pipeline described in our previous publications (Ma et al. 2018; Liang et al. 2012a; Liang et al. 2013; Liang et al. 2015b; Liu and Zhang 2019). First, head motion was estimated by frame-wise displacement (FD) in each rsfMRI scan (Power et al. 2012). Volumes with large FD (> 0.2mm) and their immediate preceding and following volumes were removed. The first 10 rsfMRI volumes were also removed to ensure steady-state magnetization. SE-rsfMRI scans with >10% volumes scrubbed were excluded from further analysis. Subsequently, the procedures of co-registration, spatial smoothing (Gaussian kernel, FWHM = 0.75mm), motion-correction, nuisance regression of 6 motion parameters (3 translational and 3 rotational parameters) and signals from the white matter and ventricles were respectively applied.

#### ME-rsfMRI data preprocessing

ME-rsfMRI data were used to separate neural and non-neural signals in rsfMRI data (Kundu et al. 2013; Kundu et al. 2012; Kundu et al. 2014). This capacity is based on the premise that neural activity-induced blood-oxygenation-level dependent (BOLD) signal change is TE dependent, as BOLD changes originate from alterations in paramagnetic deoxy-hemoglobin concentrations, which lead to changes in T*_2_ values. An innovative feature of the ME-EPI method is that each rsfMRI volume was acquired at multiple TEs, which allowed us to examine voxel-wise TE dependency of rsfMRI signal (see below).

Preprocessing of ME-rsfMRI data was similar to SE-rsfMRI data. The same motion scrubbing method described above was used to remove volumes with FD > 0.2 mm. The first 10 volumes were also removed to ensure steady-state magnetization. ME-rsfMRI scans with > 10% volumes scrubbed were excluded from further analysis. Motion parameters were then estimated using moderately smoothed EPI images (FWHM=0.75mm) acquired at the first echo. Subsequently, we combined estimated matrices of motion parameters and the affine matrix from co-registration, and applied the combined matrix to each volume in EPI. After that, images were spatially smoothed (Gaussian kernel, FWHM = 0.75mm). The time course of each voxel was detrended, and 6 motion parameters were regressed out.

To separate neural and non-neural signals, we employed a similar method established by Kundu et. al. (Kundu et al. 2013; Kundu et al. 2012; Kundu et al. 2014), including the steps of optimally combining ME data, spatial group ICA, dual-regression back reconstruction, and computation of BOLD and non-BOLD weighting for each ICA component.

For each voxel in ME-rsfMRI images, Eq. 1 describes the MR signal as a function of *TE*, where *S(TE)* is the signal amplitude at *TE*, *S*_0_ is the signal amplitude when *TE* is zero, and *T2\** is the time constant for *T2\** relaxation. For each scan, the rsfMRI signal of each brain voxel was averaged across volumes for each *TE*, and *T2\** was estimated by fitting signals at four *TEs* to Eq. 1. Then, signals acquired at four *TEs* were weighted averaged with the weight quantified using Eq. 2, where 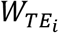 is the weight for the *i-th TE* (*TE*_*i*_). This step maximized the contrast-to-noise ratio of rsfMRI data for group-ICA.

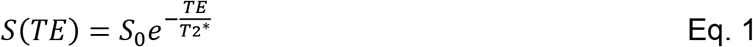

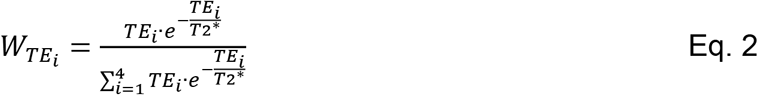

Spatial group-ICA was subsequently applied to optimally combined ME-rsfMRI data to generate group-level independent components using GIFT v3.0b (component number = 100, Fig. 1C) (Calhoun et al. 2001). Spatial maps and the corresponding time courses of all ICA components were then generated for each individual scan using dual regression back reconstruction (Calhoun et al. 2001).

To differentiate neural and non-neural components in ME-rsfMRI data, BOLD (κ) and non-BOLD (ρ) weights of each ICA component were decided. κ and ρ were pseudo-*F*-statistics that measured how likely the BOLD signal changes were due to the change of 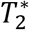 (BOLD) or *S*_0_ (non-BOLD) in Eq. [1], as BOLD signal changes were *T*2* dependent, whereas non-BOLD changes were not. Specifically, the first order approximation of Eq. [1] can be further separated into two linear sub-models:

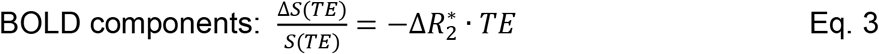

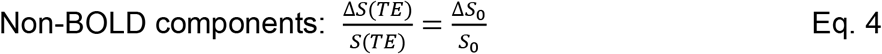

where *ΔS(TE)/S(TE)* is the signal change at a given TE. Signal changes of each voxel at different *TEs* were fit to the two sub-models, and goodness-of-fit (*F*) for Eq. 3 and 4 was calculated (i.e. 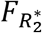 and 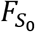), respectively. The κ and ρ weights for each ICA component were obtained by weighted averaging 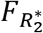 or 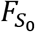of all voxels of a component using the voxel’s ICA component weight (i.e. z value) as the weighting. BOLD components had high κ, whereas non-BOLD components had high ρ. Fig. 1B shows the ρ and κ values for all ICA components, with *ρ* being sorted from the smallest to the largest. The elbow point of the sorted curve was noted as *ρ*_*elbow*_ (= 6.43). All ICA components meeting the criteria of *κ > ρ* and *ρ < ρ*_*elbow*_ were considered as BOLD components (79 in total, referred to as neural components hereafter), and all other components were recognized as non-BOLD components (21, Fig. 1C, referred to as non-neural components hereafter). Notably, these criteria were highly stringent for defining neural components. Dual regression back reconstruction was performed to derive the time courses of ICA components for each individual scan. Lastly, time courses of all non-BOLD components and the white matter signal were regressed out from optimally combined ME-rsfMRI data for each scan.

#### Generating global signal co-activation patterns

For each rsfMRI scan, the time course of each voxel was first normalized to its mean, and the global signal was calculated as the average of normalized time courses across all brain voxels, and then temporally filtered to 0.01-0.1Hz. 15% rsfMRI volumes with the highest global signal amplitude were selected for each scan, pooled together across all scans, and averaged to generate the global signal co-activation pattern (CAP). In addition, k-means clustering was applied to group these selected rsfMRI volumes based on their spatial similarity, and the mean CAP for each cluster was calculated. The same processing procedures were separately applied to SE-rsfMRI and ME-rsfMRI data.

#### Extracting spatiotemporal patterns of the global signal

To further investigate the spatiotemporal dynamics of the global signal, we extended our analysis to rsfMRI epochs surrounding local global signal peaks. Each global signal epoch was defined as rsfMRI volumes (9 s in duration) centered at a local global signal peak. Global signal epochs overlapping with each other were excluded from the analysis to ensure data independency. The volumes used for global epochs accounted for 55.8% of the total number of volumes in ME-rsfMRI data, and explained 75.0% of variance in the global signal. All global signal epochs were averaged to generate the mean spatiotemporal patterns of the global signal. This procedure was separately applied to SE-rsfMRI and ME-rsfMRI data. All global signal epochs were subjected to spatial group ICA to further extract individual brain networks involved in the global signal.

#### Surgery for calcium signal recording

rsfMRI and calcium signal signals were simultaneously recorded (n = 4) using the method previously reported (Liang et al. 2017). Before imaging, stereotaxic surgery was conducted for virus injection and optic fiber implantation. Rats (350 – 450 g) were anesthetized by intramuscular (IM) injections of ketamine (40 mg/kg) and xylazine (12 mg/kg). Buprenorphine (1.0mg/kg) was injected subcutaneously (SC) as long-term post-surgery analgesia. Dexamethasone (0.5 mg/kg) was injected SC to prevent tissue inflammation. Rats were intubated with endotracheal catheter with a fiber-optic guide (Rivera et al. 2005), placed on a stereotaxic frame (David Kopf Instruments, Tujunga, CA), and ventilated with oxygen. The animal’s heart rate and SpO2 were monitored and the body temperature was maintained at 37°C during surgery. A small craniotomy was made unilaterally over the dorsal hippocampus (dentate gyrus, 3.5 mm rostral and 2 mm lateral to bregma). AAV5.Syn.GCaMP6s (800-1000 nl, Penn Vector Core) was injected through a micropipette syringe fitted with a glass pipette tip (Hamilton Company, Reno, NV) at three depths: −2.6 mm, −2.8 mm, and −3.0 mm (~300 nl at each depth). After virus injection, a fiber optic (400 μm core, 0.48NA, 2.5mm ceramic ferrule, Thorlabs, Newton, NJ) was advanced to the injection site (depth: −2.6 mm). Five screws (0.06 inch in diameter, 0.125 inch in length, brass; McMaster-Carr, Aurora, OH) were implanted along the temporal ridge of the rat skull. Dental adhesive and dental cement (ParaBond, COLTENE, Cuyahoga Falls, OH) were applied to cover the skull and fix the implanted fiber. Rats were returned to home cages, and allowed for recovery and GCaMP expression for at least 4 weeks.

#### Simultaneous calcium-rsfMRI signal recording

A two-wavelength GCaMP fiber photometry system (Doric Lenses Inc., Quebec, Canada) was utilized for calcium signal recording (Fig. 2A) (Kim et al. 2016). GCaMP and Ca^2+^-independent fluorescent signals were alternatingly excited by a 465nm (7μW) LED and an isosbestic wavelength (405 nm, 0.75μW) LED, respectively, both modulated at 400Hz (50% duty cycle for each wavelength). Power for the two LEDs was adjusted to achieve comparable intensity of emitted fluorescent light for the two channels due to the different absorption rate of the two wavelengths in the brain tissue and different light efficiency in the optic setup. Both excitation sources were combined via a dichroic mirror in a mini-cube and coupled into a mono fiber optic patch cable (400 nm core, 0.48NA, 7m long, Doric Lenses Inc., Quebec, Canada) connected with the implanted optical fiber. Emitted fluorescent light was collected through the same patch cable, separated by another dichroic mirror in the mini-cube, coupled via a fiber launch (Thorlabs, Newton, NJ) and a 40×0.65 NA microscope objective (Olympus, Center Valley, PA), and then focused into a photomultiplier (MiniSM 30035, SensL Technologies, Somerville, MA). The converted signal was amplified (bandpass filtered at 0.3-1kHz, Dagan Corp., Minneapolis, MN) and recorded using an NI-DAQ board (10kHz sampling rate, NI USB-6211, National Instruments, Austin, TX) and custom-written LabVIEW code. TTL signals used to control the two LED modules were also recorded. We used a time-division multiplexing strategy to time-sequentially sample 465 nm and 405 nm excited fluorescent signals (Fig. 2). 405 nm signals were regressed out from 465 nm signals to correct for fluorescence changes unrelated to neuronal activity (Kim et al. 2016).

**Figure 2.**
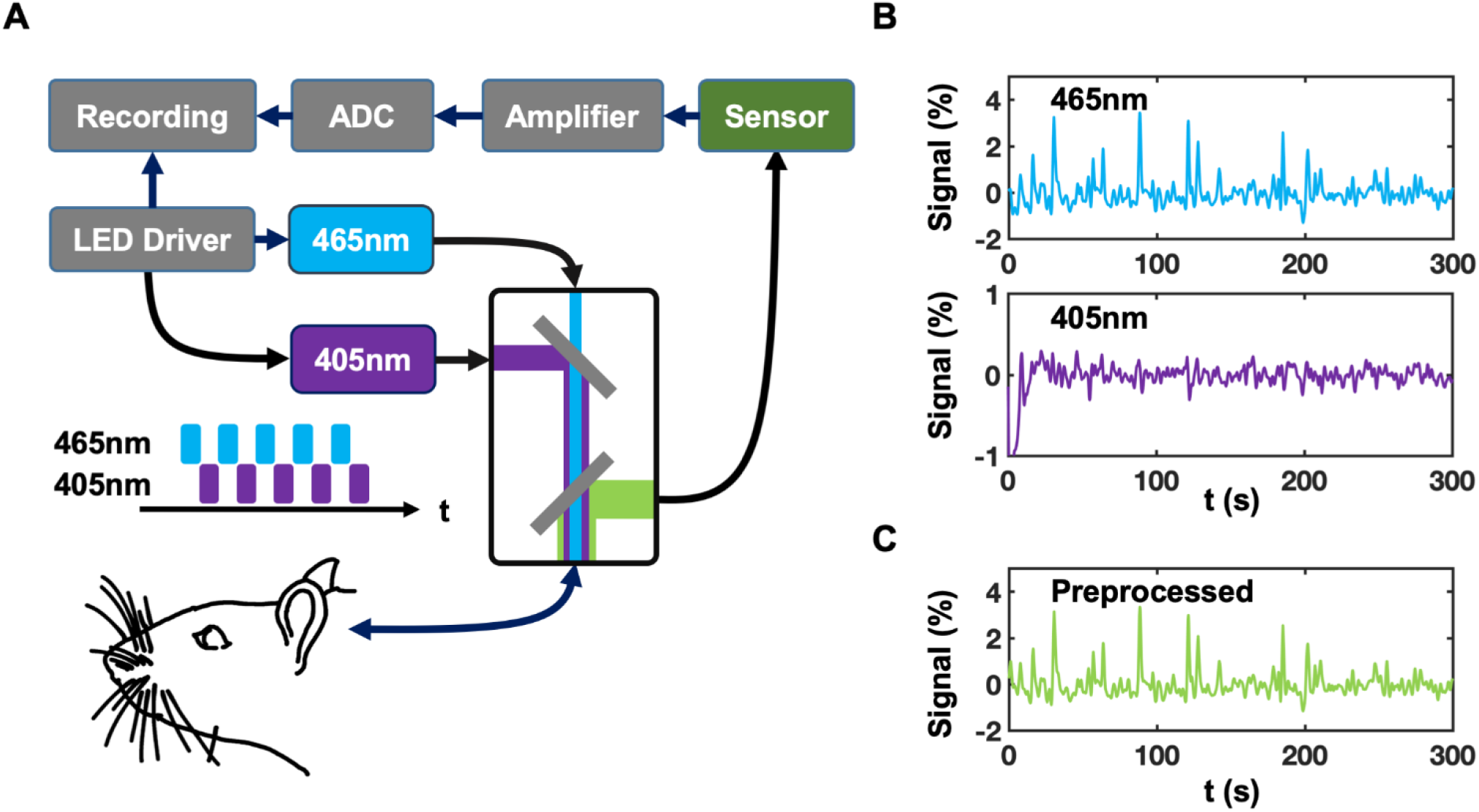
Setup of GCaMP fiber photometry and signal preprocessing. (**A**) Setup of the two-wavelength GCaMP fiber photometry system. (**B**) Signals detected with 465 nm and 405 nm excitation wavelengths, respectively. (**C**) 405 nm signals were regressed out from 465 nm signals to provide fluorescence changes pertinent to neuronal activity.

Along with GCaMP signal, SE-rsfMRI signal was simultaneously collected using the same imaging parameters mentioned above. We acquired 1-3 scans in each session. For each animal, we performed multiple sessions on separate days, which provided 20 scans in total.

## Results

In this study we systematically investigated the global brain signal in awake rats. We first examined the spatial patterns of rsfMRI volumes at global signal peaks. The neural components in the global signal spatial pattern were further investigated using ME-rsfMRI data and simultaneously recorded calcium signal.

### Co-activation patterns of the global rsfMRI signal

Fig. 3 shows the mean CAP generated by averaging 15% rsfMRI volumes with the highest global signal amplitude. This global signal CAP displayed well-organized network activities mainly involving the thalamocortical and hippocampocortical networks. In particular, regions in these two networks including the medial prefrontal, insular, anterior cingulate, retrosplenial and sensorimotor cortices, as well as medial dorsal thalamus, hippocampus and basal forebrain showed strong BOLD signal during global signal peaks. This result suggests that the global signal in awake rats might be attributed to coordinated activities in the thalamo-cortical and hippocampo-cortical networks.

**Figure 3.**
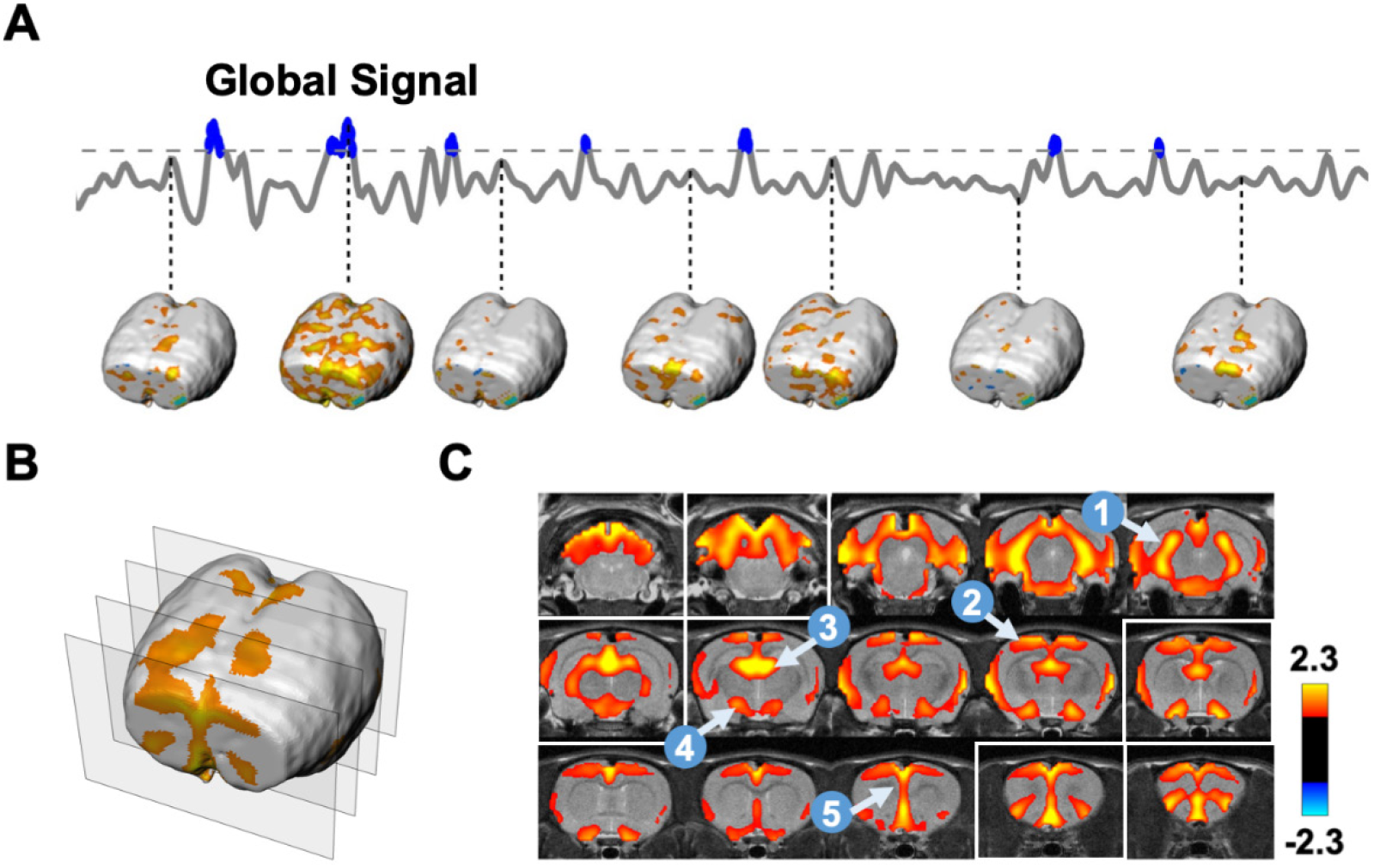
Averaged co-activation pattern (CAP) during global signal peaks in SE-rsfMRI data. (**A**) Extraction of rsfMRI volumes during global signal peaks. (**B**) 3D visualization of the averaged CAP during global signal peaks. Positions of the four slices selected were highlighted in white boxes in (**C**). (**C**) Slice-by-slice view of the global signal CAP. Five brain regions highlighted include: 1, hippocampus; 2, sensory-motor cortex; 3, medial dorsal thalamus; 4, basal forebrain; 5, prefrontal cortex.

Averaging all rsfMRI volumes selected could potentially mask distinct CAPs in the global signal. To examine this possibility, k-means clustering (k = 12) was used to group rsfMRI volumes based on their spatial similarity (Fig. 4). Fig. 4A displayed clusters of CAPs resembling the mean global signal CAP (spatial correlation coefficient = 0.72±0.10), which accounted for 79% of total rsfMRI volumes selected. In contrast, several other cluster CAPs exhibited spatial patterns quite different from the mean global signal CAP (Fig. 4B). These patterns occurred in a minor portion (21%) of rsfMRI volumes selected. None of them showed any patterns consistent with known anatomical or network structures, and therefore, might be attributed to non-neural artifacts. Taken together, our SE-rsfMRI data demonstrated that there were significant neural and non-neural contributions to the global signal, even after routine signal preprocessing.

**Figure 4.**
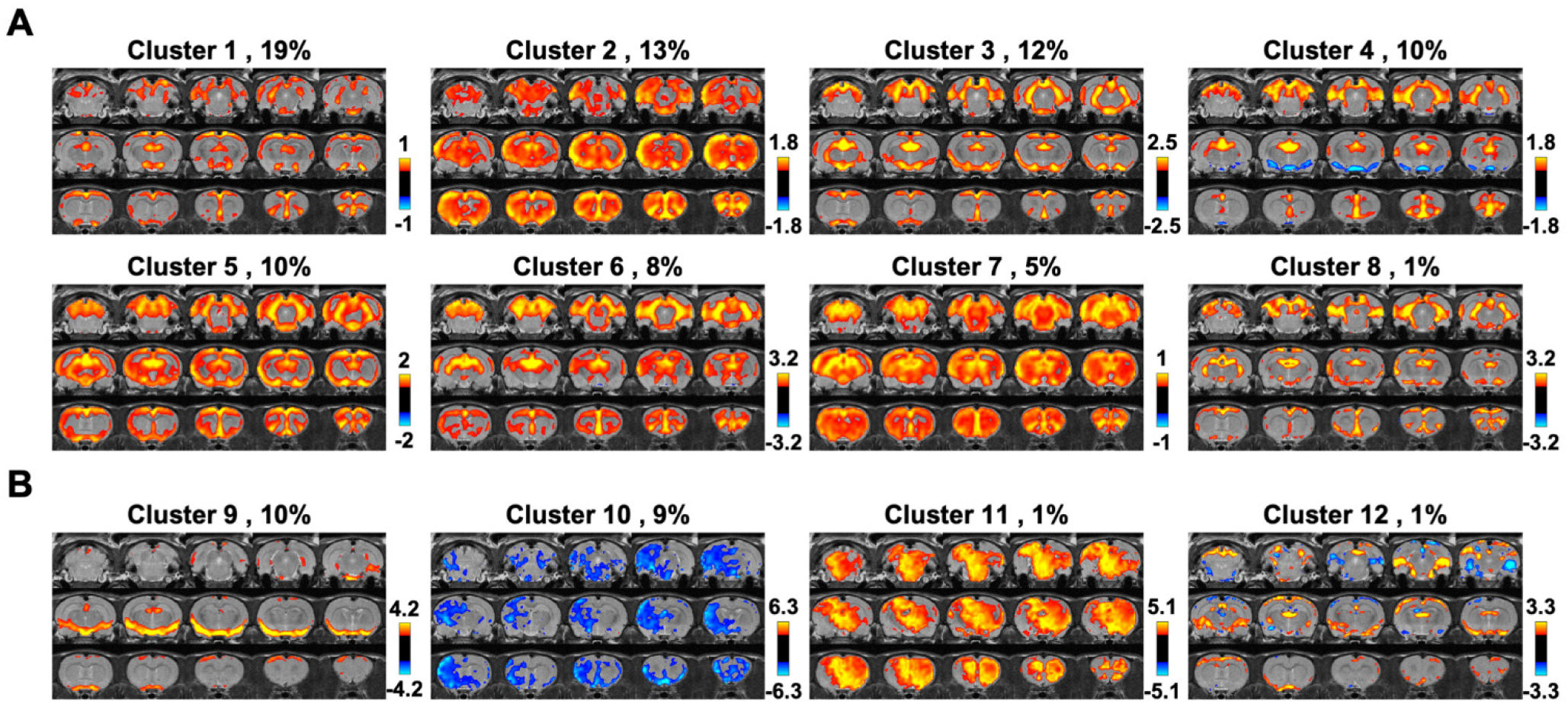
Clustering rsfMRI frames during global signal peaks. (**A**) Clusters resembling the mean global signal CAP. (**B**) Clusters exhibiting distinct CAPs. The occurrence rate is displayed next to the cluster number.

### Dissecting neural components in the global rsfMRI signal

ME-rsfMRI was employed to further differentiate neural and non-neural components in the global rsfMRI signal. Fig. 1C shows both neural and non-neural components generated using ME-rsfMRI data. rsfMRI signals from all non-neural components were regressed out from ME-rsfMRI data, generating a de-noised rsfMRI dataset. The mean CAP of all global signal peaks was then calculated using the same method described above. Our data showed that the mean global signal pattern in de-noised ME-rsfMRI data (Fig. 5) was highly consistent with the mean global signal CAP obtained from SE-rsfMRI data (Fig. 3C), including activated regions in the thalamo-cortical and hippocampo-cortical networks, as well as the basal forebrain. Consistent results between SE- and ME-rsfMRI data further validated that the global signal in awake rats was linked to coordinated neural activities in these networks.

**Figure 5.**
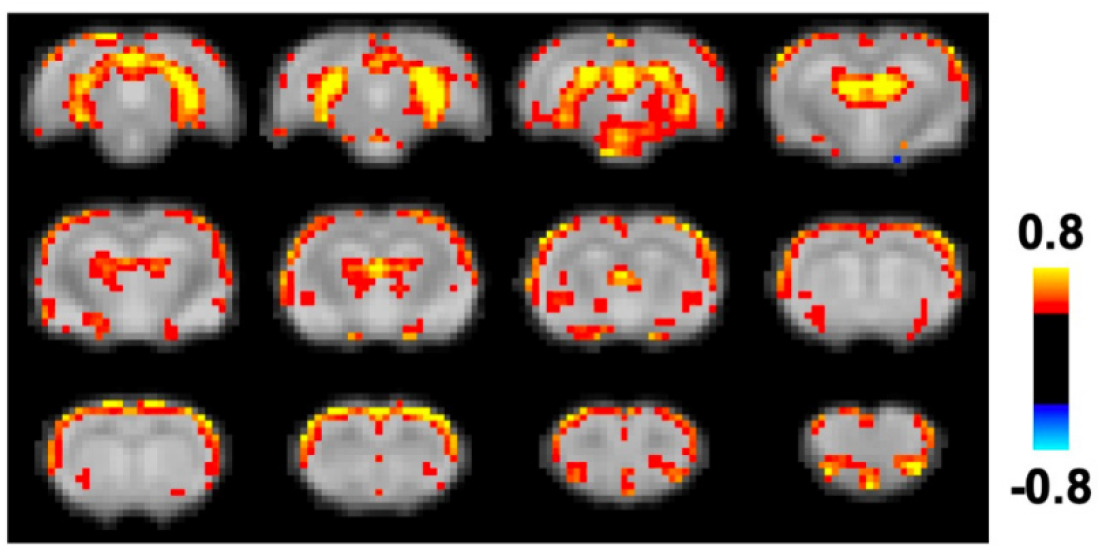
Co-activation pattern during global signal peaks in de-noised ME-rsfMRI data.

### Spatiotemporal dynamics of the global signal

To examine the spatiotemporal dynamics of the global signal, we analyzed rsfMRI epochs surrounding local peaks of the global signal (9 s per epoch, Fig. 6A). Figs. 6B & 6C show the spatiotemporal patterns averaged across all global signal epochs for SE-rsfMRI and ME-rsfMRI data, respectively. As expected, the central volumes in both SE and ME-rsfMRI data were highly consistent with the mean global signal CAPs shown in Figs. 3C and 5, respectively, with strong activity in the thalamo-cortical and hippocampo-cortical networks. The involvement of these two functional networks in global signal dynamics was further validated by spatial group-ICA applied to all global signal epochs in the ME-rsfMRI dataset, which provided two prominent ICA components representing the thalamo-cortical network and hippocampo-cortical network, respectively (Fig. 7).

**Figure 6.**
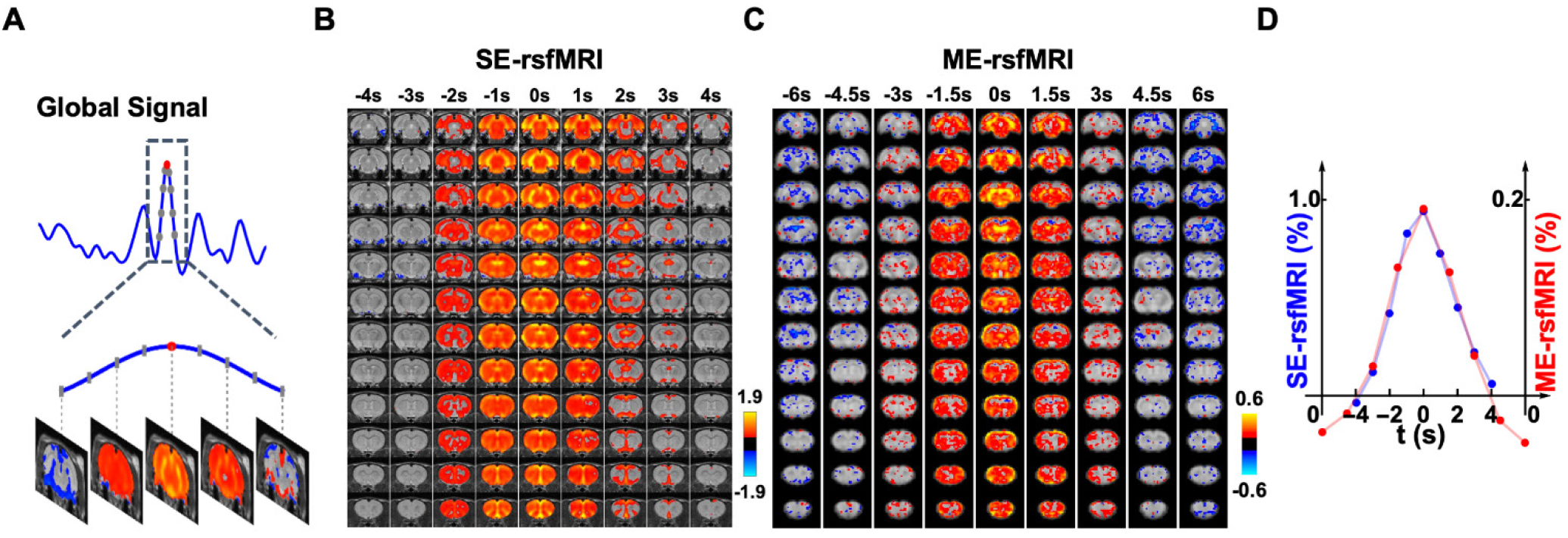
Spatiotemporal dynamics of the global signal. (**A**) An example of a global signal epoch including 8 rsfMRI volumes (gray dots) surrounding a local peak (red dot). (**B**) Averaged spatiotemporal pattern of global signal epochs in SE-rsfMRI data. Each column represents the averaged spatial pattern of global signal epochs at a time point. (**C**) Averaged spatiotemporal pattern of global signal epochs in ME-rsfMRI data. (**D**) Frame-by-frame global signal amplitude averaged across all epochs in SE and ME-rsfMRI data, respectively.

**Figure 7.**
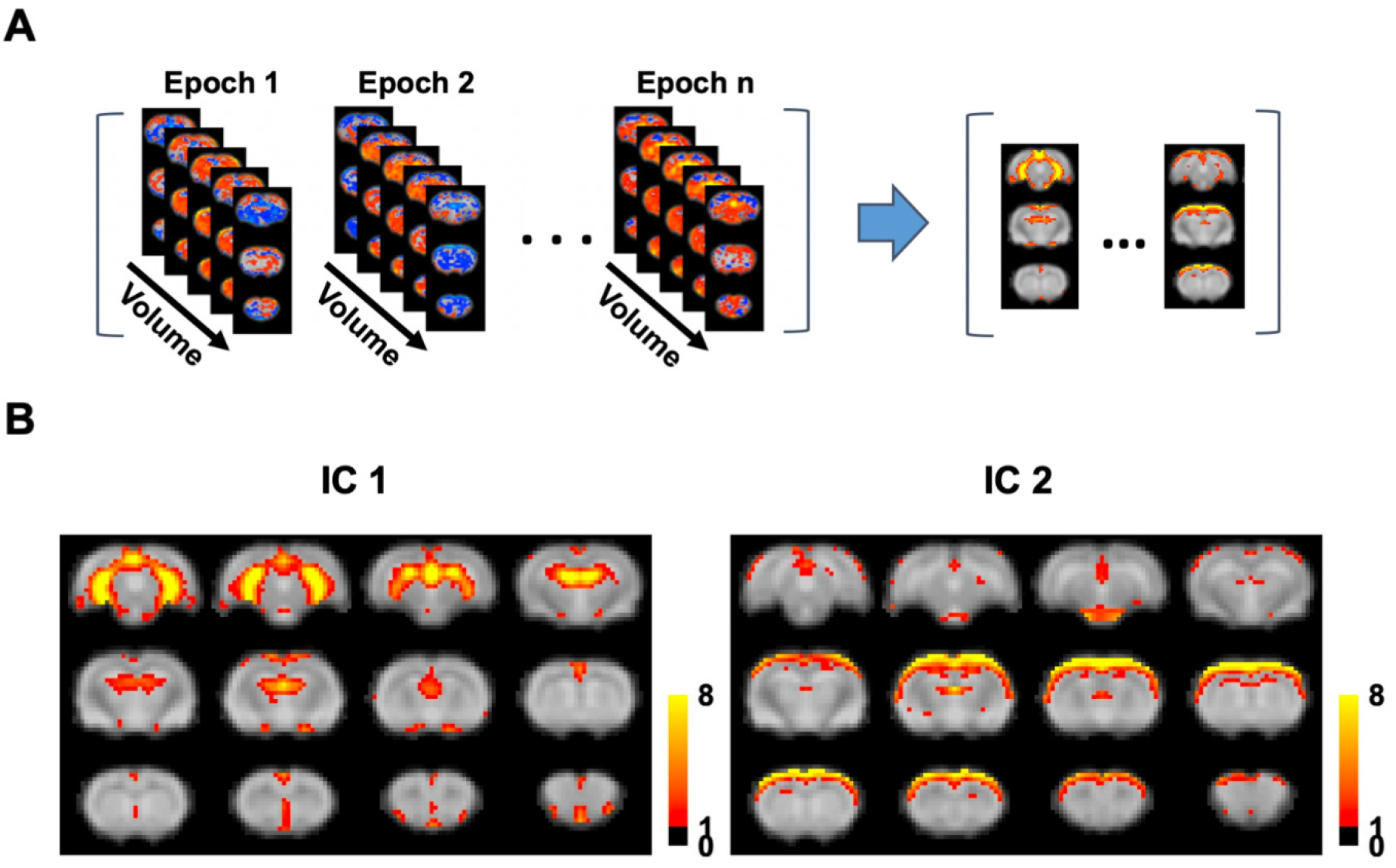
Major functional networks involved in the global signal. (**A**) Spatial group-ICA applied to all global signal epochs obtained using ME-rsfMRI data. (**B**) Two major functional networks derived from group ICA.

Fig. 6D shows frame-by-frame global signal amplitude averaged across all global signal epochs for SE- and ME-rsfMRI data, respectively. Results from both datasets displayed almost identical global signal temporal profiles, which started ~4 s before the peak, sustained for ~2-3 s and then returned to baseline, with a total period of ~8s.

### Correspondence between the global rsfMRI signal and neuronal spikes in the hippocampus

To further validate that the global signal involved neural activity in the hippocampo-cortical network, we concurrently recorded the spiking activity in the dentate gyrus using GCaMP-based fiber photometry and rsfMRI data in awake rats. The correspondence between the GCaMP and global rsfMRI signals was quantified by estimating the portion of global signal peaks following GCaMP peaks within the time window of 2-6 sec (i.e. hemodynamic delay).

To test the statistical significance of the correspondence between the calcium and global signals, we used a permutation test by randomizing the position of global signal peaks in each scan and re-calculated the co-occurrence rate between the two signals. The permutation process was repeated 1000 times to obtain the null distribution of the co-occurrence rate. This statistical test demonstrated significant correspondence between neural spikes in the hippocampus and global rsfMRI signal peaks (p = 0.016) (Fig. 8), again confirming that the global rsfMRI signal was associated with hippocampal neural activity.

**Figure 8.**
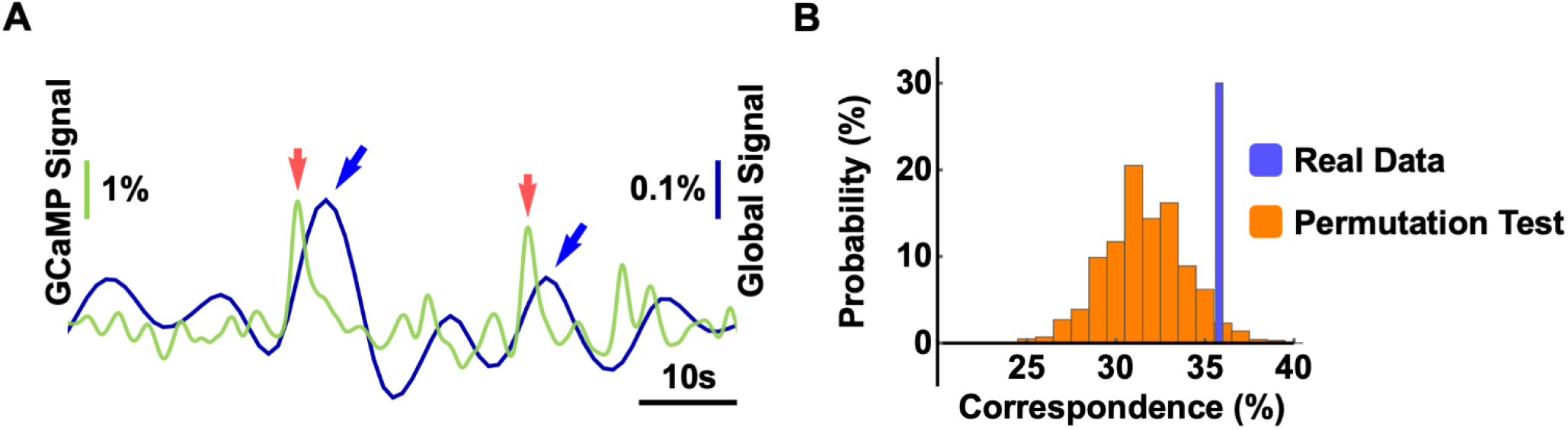
Correspondence between the global rsfMRI signal and neuronal spikes in the hippocampus. (A) Representative time courses of the global rsfMRI signal (blue) and calcium signal (green). Global signal peaks (blue arrows) and neural spikes (red arrows) in the hippocampus were strongly coupled within 2-6 sec of hemodynamic delay. (B) Permutation test of the correspondence between the global signal and GCaMP peaks (p = 0.016, 1000 permutations).

Notably, our data showed that ~35% global signal peaks were related to hippocampal spikes, but not all. To further examine this issue, we obtained the BOLD pattern of GCaMP-corresponded global signal peaks (35% volumes, Fig. 9A) and those not corresponding to GCaMP peaks (65% volumes, Fig. 9B). The data demonstrate that the spatial map of GCaMP-corresponded global signal peaks was highly consistent with the global signal pattern identified (Fig. 3), whereas the BOLD map of global signal peaks that did not correspond to hippocampal spikes displayed a less similar pattern. It needs to be noted that the latter could still be contributed by neural activity in other regions in the thalamocortical and/or hippocampal networks. Indeed, some cortical activity was observed in this pattern, but the activity in the hippocampus, medial prefrontal cortex and thalamus was largely absent. This pattern also appears noisier, consistent with our SE-rsfMRI data which showed that ~21% global signal peaks were from non-neural sources (Fig. 4).

**Figure 9.**
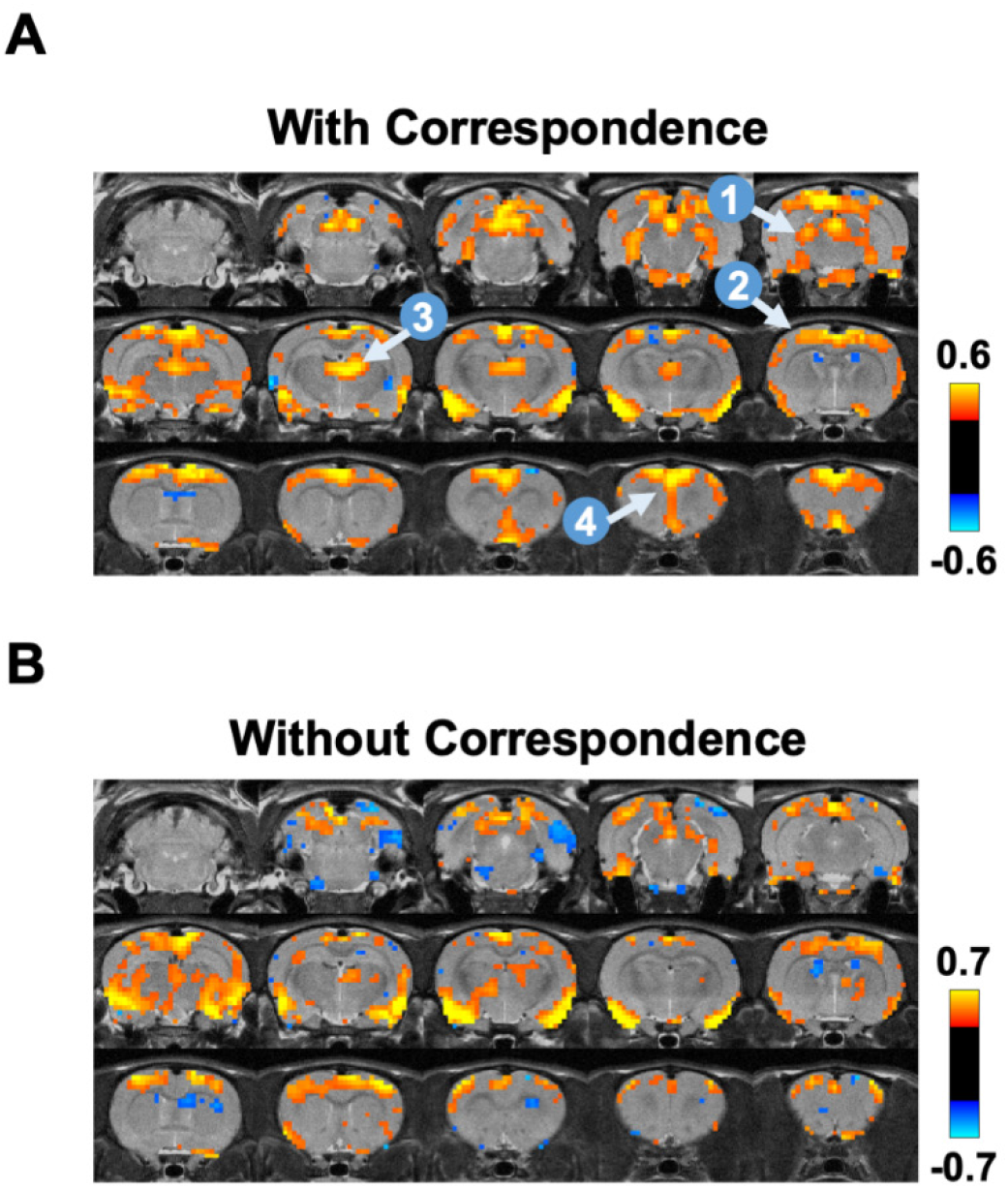
BOLD co-activation patterns during global signal peaks with (A) and without (B) corresponding GCaMP peaks in the hippocampus. Four brain regions highlighted include: 1, hippocampus; 2, sensory-motor cortex; 3, medial dorsal thalamus; 4, prefrontal cortex.

### Impact of head motion and respiration

Head motion and changes of respiration rate can significantly impact the whole-brain rsfMRI data (Kalthoff et al. 2011). To rule out the potential impact of these factors on the global signal pattern, we compared the FD and respiration rate for rsfMRI frames in global signal epochs and those not in global signal epochs in ME-rsfMRI data (Fig. 10). No significant difference was observed in head motion (p = 0.61, t = 0.52, df = 84) or respiration rate (p = 0.86, t = 0.17, df = 84), suggesting that neither head motion nor respiration had a dominant effect on the global signal pattern. In addition, the averaged motion level was on the order of 50 μm, far less than the voxel size (500 × 500 × 1000μm^3^), suggesting that that the overall motion level was low in our data.

**Figure 10.**
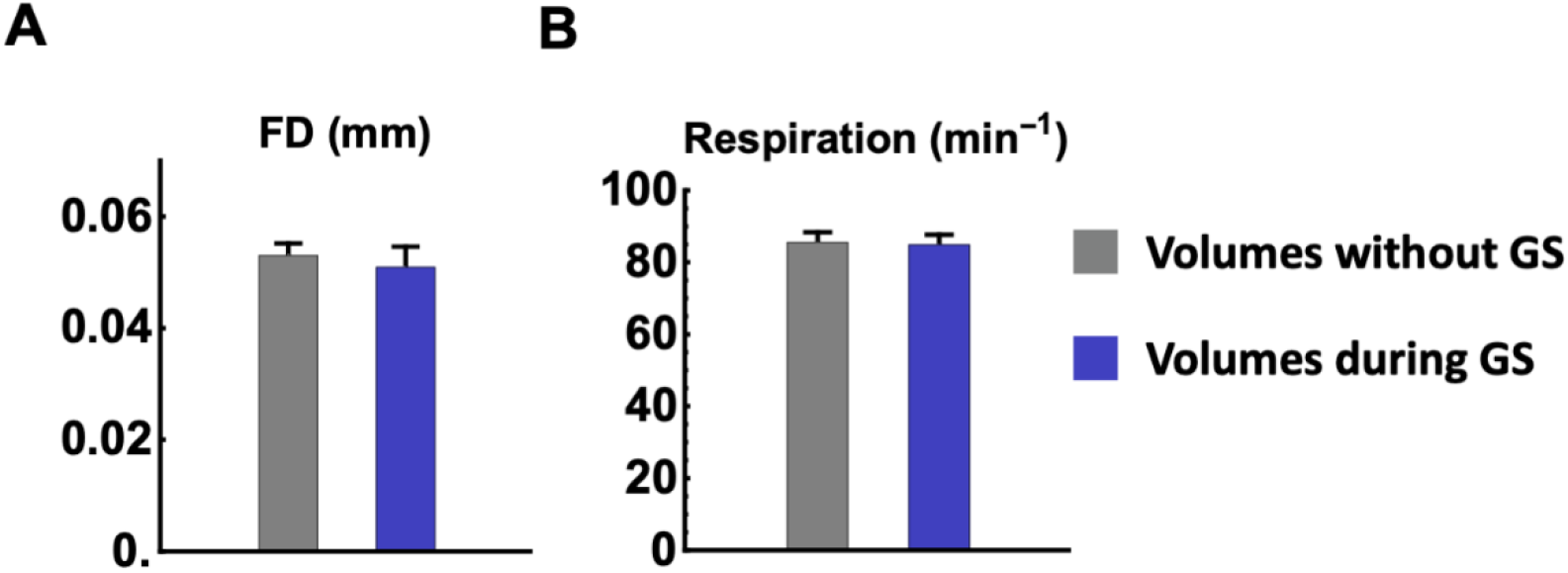
Effects of head motion and respiration. (**A**) Framewise displacement (FD) and (**B**) respiration rate for rsfMRI frames in global signal epochs (blue bars) and rsfMRI frames not in global signal epochs (gray bars), respectively. No significant difference was found in head motion level (*p* = 0.61) or respiration (*p* = 0.86).

## Discussion

In the present study we systematically investigated the global brain signal in awake rodents. Using SE-rsfMRI, we found that the global signal might involve neural activity in the hippocampo-cortical and thalamo-cortical networks. This finding was corroborated by ME-rsfMRI data after non-neural components were extensively removed. To further validate the neural origin of the global signal pattern, we simultaneously recorded calcium and rsfMRI data in awake rats, and found significant correspondence between neural spikes in the hippocampus and global signal peaks. Collectively, these data have filled the knowledge gap of the global brain signal in awake rodents, and provided strong evidence supporting its neural basis.

### Non-neural or neural?

It is well recognized that the global brain signal contains significant non-neural components including physiologic fluctuations, head motion and scanner instability (Liu et al. 2017; Nalci et al. 2017). This nature makes global signal regression an effective method to remove artifacts in rsfMRI data. However, doubt has been casted on the validity of this preprocessing procedure as it has been shown that global signal regression mathematically mandates negative correlations and might cause artifactual anticorrelated RSFC (Murphy et al. 2009). Studies have also shown that, besides non-neural components, the global signal has neural basis (Scholvinck et al. 2010), further suggesting that global signal regression can remove meaningful neural activity/connectivity, and thus bias RSFC quantification. At the functional level, the global signal is tightly related to vigilance in human subjects. Wong and colleagues found that the global signal amplitude was correlated to the vigilance level measured using EEG (Nalci et al. 2017; Wong et al. 2012, 2013), and could be modulated using caffeine (Rack-Gomer and Liu 2012). Recent studies further elucidated specific whole-brain CAPs during arousal level changes that were coupled to the global signal (Liu et al. 2018; Chang et al. 2016a). Moreover, the global brain signal was found to be altered in patients with schizophrenia, suggesting that it may be involved in neuropathophysiology of psychiatric disorders (Yang et al. 2017; Yang et al. 2014). Taken together, there is compelling evidence supporting that the global brain signal might contain significant neural components in humans and anesthetized animals.

Consistent with this notion, our data demonstrate that the global brain signal in awake rodents has important neural basis, which might involve coordinated neural activity in the hippocampo-cortical and thalamo-cortical networks. First, we observed strong co-activations in highly structured networks during global signal peaks in SE-rsfMRI data. These network structures consistently maintained even if we clustered rsfMRI frames based on their spatial similarity. Second, the same network activity was also observed in ME-rsfMRI after extensively removing non-neural components. Third, the global signal peaks corresponded to simultaneously measured neural spiking activity in the hippocampus. Taken together, these data have provided strong evidence supporting the neural basis of the global rsfMRI signal in awake rodents.

### Functional networks involved in the global brain signal

We found two prominent functional networks co-activated during global signal peaks – hippocampo-cortical and thalamo-cortical networks. Strong interactions between the hippocampus and the cortex have been reported in both rodent and monkey studies (Chan et al. 2017; Logothetis et al. 2012). Such interactions are often linked to brain-wide activity and might play an important role in brain function. For instance, optogenetic stimulation of the dentate gyrus was shown to enhance brain-wide functional connectivity at the resting state in rats (Chan et al. 2017). In monkeys, sharp-wave ripples in the hippocampus were tightly coupled to increased activity in virtually all cortical regions, which hinted the relationship of this hippocampo-cortical co-activation and the global brain signal given the large portion of brain volume involved (Logothetis et al. 2012; Ramirez-Villegas et al. 2015). Functionally, the activity of the hippocampo-cortical network was believed to relate to the consolidation of hippocampus-dependent memory (Ramirez-Villegas et al. 2015). Like the hippocampo-cortical network, the thalamo-cortical network was also found to be involved in the global brain signal. A recent rodent study reported that low-frequency optogenetic stimulations of the ventral posteromedial thalamus led to brain-wide neural activity via both mono and multi-synaptic projections to cortical regions, alluding the potential contribution of the thalamocortical network to the global signal (Leong et al. 2016). Although specific regions involved may differ, these studies all support that activities in the hippocampo-cortical and thalamo-cortical networks play a major role in the global brain signal in rodents.

In addition to the two major functional networks, we observed remarkable involvement of the basal forebrain in the global brain signal. Correspondingly, recent human (Liu et al. 2018) and monkey (Turchi et al. 2018) studies demonstrated that the basal forebrain drove the brain-wide cortical activities, especially in sensory-motor areas, and regulated the global brain signal. These results collectively indicate that the global brain signal involve activities from both large-scale neural networks and small subcortical structures.

### Cross-species translation

There is compelling evidence demonstrating that the global brain signal is tightly related to the arousal level in both humans (Wong et al. 2013; Liu et al. 2018) and monkeys (Turchi et al. 2018; Chang et al. 2016a). It has been further suggested that the global brain signal and arousal were regulated by the basal forebrain (Turchi et al. 2018). Whether these results can be translated to rodents is an interesting topic to investigate and certainly warrants more detailed studies in the future. However, some differences between these species that might affect the translation of the global brain signal need to be acknowledged. In particular, in rodents two thirds of the brain volume are subcortical regions and one third is the cortex. This composition is in remarkable contrast to primates/humans, in which cortical regions occupy the vast majority of the brain volume. Given that the global brain signal is defined by the averaged activity across the whole brain, it would have very different contributions from the cortex versus subcortex between rodents and primates/humans. Therefore, it should not be assumed that our findings in the global brain signal can be directly translated to other species.

### Potential pitfalls and limitations

There are a number of physiological factors that can affect rsfMRI signal and can potentially impact the global rsfMRI pattern, such as respiratory volume variability, vasomotion, brain pulsation, blood pressure, just to name a few. Although our study has been carefully designed to rule out these possibilities, including using stringent motion control, regressing out signals from the white matter and ventricles, using ME-EPI to separate non-neural components and simultaneously recording GCaMP signal in the hippocampus, caution still needs to be taken for the interpretation of the results. Specifically, it is difficult to conclude that the global signal pattern revealed is completely free from non-neural artifacts. For instance, some of the physiological factors are directly related to vascular changes, which also contribute to T2* changes, and thus cannot be ruled out by ME-EPI. In addition, signals from large veins can contribute to the global rsfMRI signal. Nonetheless, given that the consistency of multiple measurements (i.e. SE-EPI, ME-EPI and GCaMP signal), it is unlikely physiologic artifacts dominated the global brain signal pattern.

Although ME-rsfMRI has the advantage of differentiating neural and non-neural components, this method cannot guarantee each ICA component was completely neural or non-neural. In fact, all ICA components showed non-zero κ and ρ values, suggesting that they were more or less mixed with neural and non-neural constituents. In this study, we used stringent criteria to select neural components. Therefore, it is likely that some non-neural ICA components identified contained some neural elements. Nonetheless, the purpose of utilizing ME-EPI in this study is to extensively regress out non-neural components, in order to validate the results found in SE-rsfMRI data. Indeed, 28 regressors were used for de-noising ME-rsfMRI data, compared to only seven regressors used in preprocessing SE-rsfMRI data. Even with such stringent criteria, consistent results were obtained between SE-rsfMRI and ME-rsfMRI data, confirming that activities in the hippocampo-cortical and thalamo-cortical networks indeed represent neural contributions to the global signal in awake rats.

The present study focused on revealing the spatial pattern of brain activity when global signal was high. An equally important issue is to understand the temporal dynamics of the global signal, and a number of advanced analysis method has been applied. For instance, a recent study investigated the relationship between separate co-activation patterns and the phase of global signal (Gutierrez-Barragan et al. 2019). This issue needs to be specifically studied in the future.

## Conclusion

We systematically investigated the global rsfMRI signal in the awake rat brain. Our data suggest that the hippocampo-cortical and thalamo-cortical networks play a major role in the neural basis of the global signal. These results have filled our knowledge gap of the global brain signal in awake rodents and provided important insight into its neural substrate. The present study can further facilitate comparative studies investigating the generalized function the global signal may have across rodents and humans.

## Acknowledgments

The present study was partially supported by National Institute of Neurological Disorders and Stroke (R01NS085200, PI: Nanyin Zhang, PhD) and National Institute of Mental Health (R01MH098003 and RF1MH114224, PI: Nanyin Zhang, PhD).

## Compliance with Ethical Standards

### Conflict of interest

none.

### Research involving Human Participants and/or Animals

The research involved animals. All procedures were conducted in accordance with approved protocols from the Institutional Animal Care and Use Committee (IACUC) of the Pennsylvania State University.

### Informed consent

N/A

### Funding

The present study was supported by National Institute of Neurological Disorders and Stroke (R01NS085200, PI: Nanyin Zhang, PhD) and National Institute of Mental Health (R01MH098003 and RF1MH114224, PI: Nanyin Zhang, PhD).

